# Small molecule inhibition of CPSF3 impacts R-loop distribution and abundance

**DOI:** 10.1101/2025.05.07.652284

**Authors:** Cristina R. Hofman, Victor Tse, Jiaxin Hu, David R. Corey

## Abstract

R-loops are three-stranded nucleic acid structures consisting of an RNA/DNA hybrid and a displaced strand of DNA. These structures have been implicated in a variety of regulatory cellular processes. Their untimed or excess accumulation, however, can cause genomic instability and induce DNA damage. Most R-loops form co-transcriptionally when the nascent transcript reanneals to unwound DNA duplex. Changes in the rate of transcription have the potential to impact R-loop formation, and compounds that modulate R-loop formation would be useful molecular tools and therapeutic leads. Cleavage and Polyadenylation Specific Factor 3 (CPSF3) recognizes the pre-mRNA 3’ cleavage site, cleaves the transcript prior to polyadenylation, and has been linked to R-loop formation. Inhibition of CPSF3 has been found to induce transcriptional readthrough and cell proliferation defects. A previous report suggested that inhibition of CPSF3 with a small molecule causes a global increase in R-loop abundance. Here we test the impact of YT-II-100, a novel inhibitor of CPSF3. We find that addition of YT-II-100 increases global R-loop formation but does not change R-loop formation at specific genes that are normally used as positive controls for R-loop formation. We performed parallel assays using previously reported compound JTE-607 and observed similar results. Our data emphasize the need for cautious interpretation of experiments using JTE-607 and YT-II-100. There may be different mechanisms of R-loop formation depending on gene loci, with the control of R-loop formation at some genes diverging from the regulation of global R-loop formation.

## Introduction

Regulation and modulation of biologically relevant structures is important for controlling endogenous gene expression, molecular tools, and therapeutic development. One example, R-loops, are three stranded nucleic acid structures composed of an RNA/DNA hybrid and a displaced strand of DNA^1^. These structures largely form in *cis*, co-transcriptionally, but can also form in *trans*. Endogenous regulation of R-loops is controlled by a variety of enzymes, including ribonuclease H (RNase H)^2^, topoisomerases^3^, and helicases^4^. Most commonly, R-loops are characterized for their role in immunoglobulin class switch recombination^5–8^ but also have roles in transcriptional regulation and maintenance of genome stability^9–13^. While important for biological regulation, excess and untimed R-loops have also been associated with genomic instability and multiple disease states^14–16^.

Cleavage and polyadenylation of newly transcribed pre-mRNA is necessary for the maturation of mRNAs. The Cleavage and Polyadenylation Specific Factor (CPSF) protein complex recognizes the AAUAAA sequence and downstream element on the nascent RNA^17,18^. After recruitment of other proteins, there is a two-step catalytic reaction that occurs, where first the transcript is cleaved and second the poly(A) tail is added to the 3’ end of the cleavage product. One component of this complex is cleavage and polyadenylation specificity factor 3 (CPSF3 or CPSF73). CPSF3 is a member of the metallo-β-lactamase superfamily of zinc-dependent hydrolases^19^ and has endonuclease activity necessary to catalyze efficient cleavage and polyadenylation^19,20^.

In 2020, Beckwith and colleagues investigated the molecular target and mechanism of action of JTE-607 (N-[3,5-Dichloro-2-hydroxy-4-[2-(4-methyl-1-piperazinyl)ethoxy]benzoyl]-L-phenylalanine ethyl ester dihydrochloride) (Fig. 2A)^21^, a small molecule antiproliferative that has progressed to early-stage clinical trials even though its mechanism of action was unknown. Using quantitative chemical proteomics, they identified CPSF3 as a molecular target for JTE-607. Physical association was supported by activity assays, binding assays, and 2.3 Å co-crystal structure.

JTE-607 inhibits the nuclease activity of CPSF3, inducing transcription read-through. They hypothesized that reduced transcript cleavage and increased transcript readthrough contributes to the observed cell death. Enhancing transcription readthrough might also increase R-loop formation. Semi-quantitative microscopy assays suggested that the addition of JTE-607 increases global formation of R-loops^21^. More recently, a study in an ovarian cancer model suggested that JTE-607 treatment has no impact on R-loop abundance^22^. The contrasting findings from these two studies suggest a complexity to the downstream effects of CPSF3 inhibition and a need for additional exploration of the link between CPSF3 and R-loop formation.

In 2024, Nijhawan and coworkers examined the molecular mechanism and targets of anticancer benzoxaboroles^23^. They used a forward genetic screen that employs iHCT116 cells, a barcoded population of cells that allows control over the frequency of mutation. These assays showed that compound YT-II-100 (3-(1,3-dihydro-1-hydroxy-2,1-benzoxaborol-7-yl)-N-(4’-acetyl-1,1’-biphenyl-3-yl) propanamide) (Fig. 1A) was antiproliferative and that mutations in CPSF3 led to resistance. The mechanism of action and binding of YT-II-100 relative to CPSF3 was confirmed by proliferation assays, binding assays, mRNA cleavage assays, and a 1.7Å co-crystal structure. Overlay of the binding of JTE-607 and YT-II-100 showed that the compounds bound within the same active site pocket and had some overlap, while CPSF3 conformational changes differed to accommodate the respective compounds. The similarities between JTE-607 and YT-II-100 with regards to activity, association with CPSF3, the structural details of association with CPSF3, and the previous observation of a potential link between R-loop abundance and JTE-607 led us to hypothesize that YT-II-100 might also have a potential role in controlling R-loop formation.

**Figure 1.**
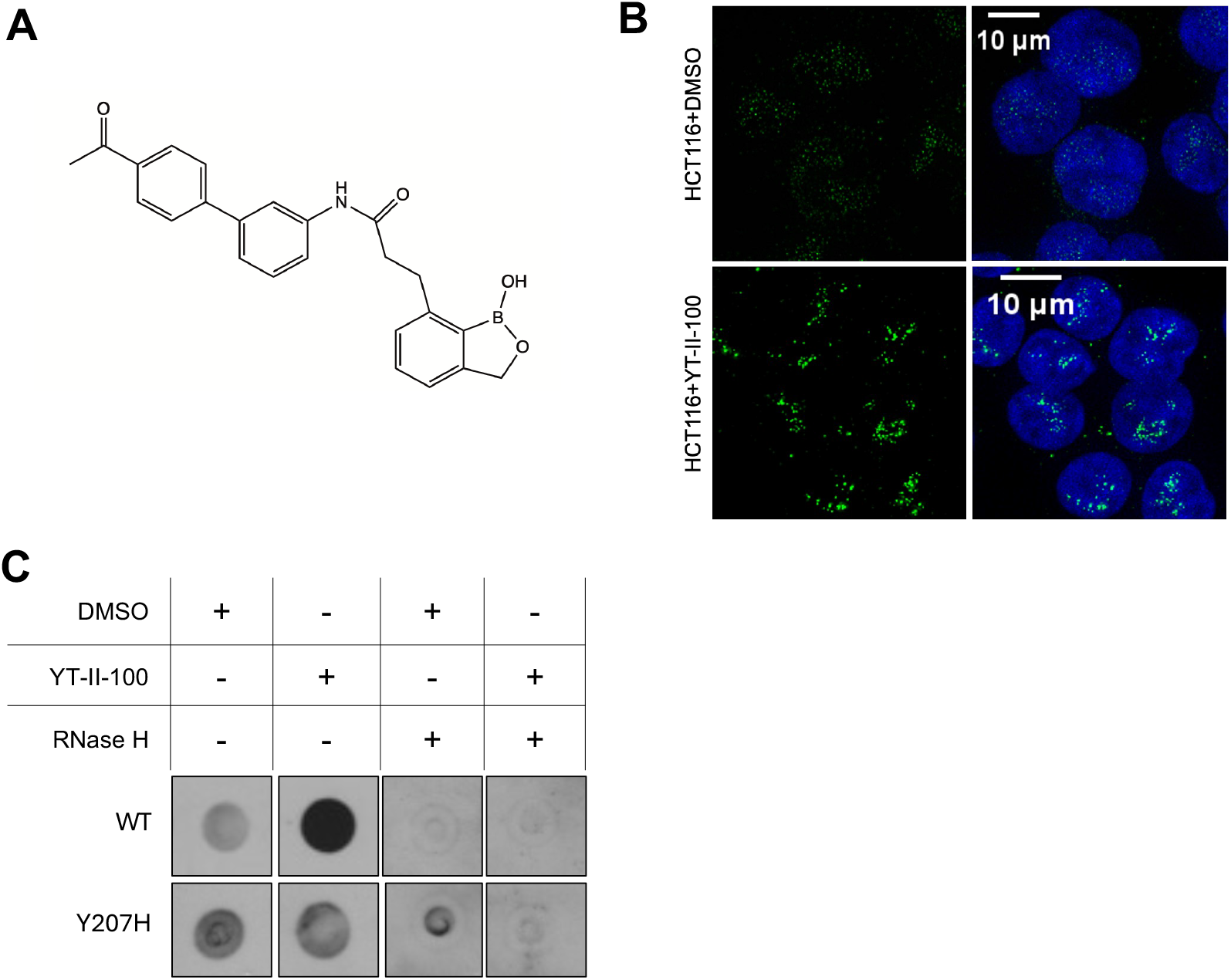
Inhibition of CPSF3 with YT-II-100 leads to a global increase in R-loops A) Chemical structure of YT-II-100. B) Fluorescence microscopy images of wild-type cells staining for RNA/DNA hybrids (green) and DAPI/cell nuclei (blue) following 6 hours of DMSO or YT-II-100 treatment. C) Dot blot staining for RNA/DNA hybrids of wild-type and Y207H HCT116 cells treated with DMSO or YT-II-100 for 6 hours +/-RNase H treatment (N=3).

Here, we use three different assays (microscopy, dot blot, and DNA/RNA immunoprecipitation) in conjunction with a cell line that expresses a mutant CPSF3 that is resistant to YT-II-100. We demonstrate using these complementary assays that YT-II-100 can induce global increases in R-loop abundance. However, while we observe global changes, we do not observe significant changes at well-characterized sites that are known to form detectable R-loops. Parallel assays using JTE-607 produce similar results. Our findings suggest that, while small molecule mediated inhibition of CPSF3 does impact global R-loop abundance, regulation of R-loop formation of some well-known “hallmark” genes is less predictable. The mechanism of this regulation is multifaceted and at least partially dependent on genomic context.

## Results

### Global R-loop abundance is upregulated following CPSF3 inhibition

The S9.6 antibody recognizes RNA/DNA hybrids^24–27^, making it a valuable reagent to monitor R-loops. To determine whether inhibition of CPSF3 by YT-II-100 (**Fig. 1A)** impacts global R-loop abundance in HCT116 colorectal cancer-derived cells, we first used immunofluorescence staining with the S9.6 antibody to recognize R-loops, to detect global levels of R-loops.

While the S9.6 antibody is specific for RNA/DNA hybrids, it does have some affinity for dsRNA and dsDNA^24^. Further, since R-loops mainly form co-transcriptionally, they are expected to be localized primarily in the nucleus. Therefore, to address the potential for non-specific S9.6 signal and increase the ability to unambiguously detect R-loops in cell nuclei, the cytoplasm of cells was lysed following YT-II-100 or DMSO treatment, and the isolated nuclei were used for immunofluorescence staining. We observed minimal S9.6 signal in cells treated with DMSO (**Fig. 2B**). Following treatment with YT-II-100, we observed enhanced punctate nuclear staining, consistent with an increase in R-loop detection. These data are consistent with previous data using JTE-607^21^ and the conclusion that YT-II-100 also increases global R-loop abundance.

**Figure 2.**
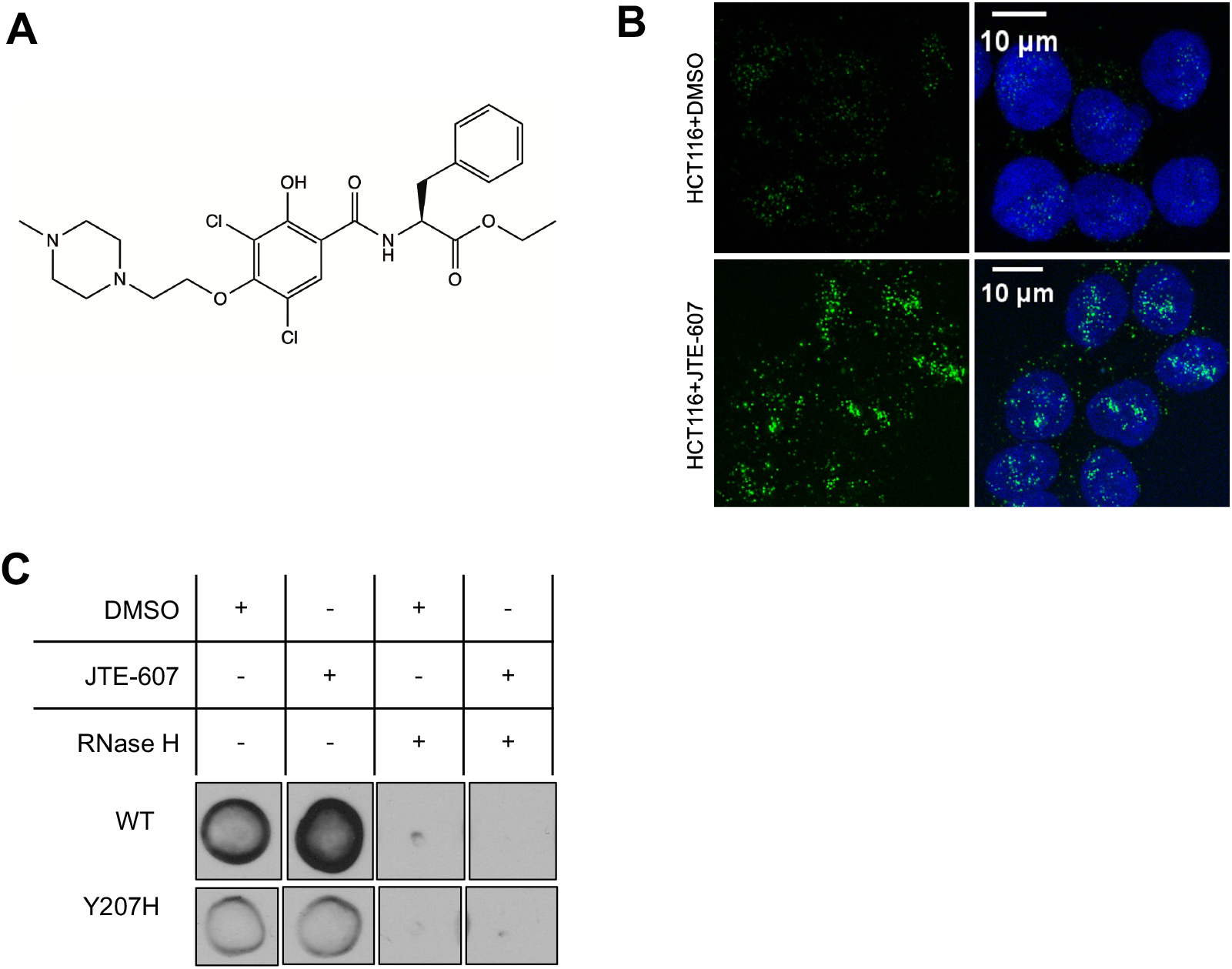
Inhibition of CPSF3 with JTE-607 leads to a global increase in R-loops A) Chemical structure of JTE-607 B) Fluorescence microscopy images of wild-type cells staining for RNA/DNA hybrid (green) and DAPI/cell nuclei (blue) following 6 hours of DMSO or JTE-607 treatment. C) Dot blot probing for RNA/DNA hybrids of wild-type and Y207H HCT116 cells treated with DMSO or JTE-607 for 6 hours +/-RNase H treatment (N=3).

We then used dot blot analysis^28^ as a complementary test for the effect of YT-II-100 on R-loops. This assay also uses the S9.6 antibody but monitors R-loop formation with a sample of isolated genomic DNA. As a control, a portion of each sample was pre-treated with RNase H, a ribonuclease that specifically degrades the RNA portion of RNA/DNA hybrids^2,29–31^. As an additional control, we used a mutant HCT116 cell line, HCT116 Y207H, which contains a point mutation in the interface between the metallo-β-lactamase and β-CASP domains of CPSF3 that abrogates YT-II-100 binding^23^. This cell line allows us to distinguish on-target effects from off-target effects.

In the wild-type cells, we observed an increase in S9.6 signal following treatment with YT-II-100 but not in samples treated with RNase H nor in samples obtained from mutant Y207H cells (**Fig. 1C**). Sensitivity to RNase H supports the conclusion that the signal was due to detection of RNA/DNA hybrids. Taken together with sensitivity to the Y207H mutation, the immunofluorescence and dot blot analysis provide strong evidence that YT-II-100 affects global R-loop abundance by a mechanism involving CPSF3.

We then evaluated a second inhibitor of CPSF3, JTE-607 (**Fig. 2A**), to further evaluate the generality of R-loop regulation by small molecules. As observed for YT-II-100, we observed that the addition of JTE-607 also increased total R-loop detection in HCT116 cells by both dot blot and microscopy assays (**Fig 2BC**). While the CPSF3 Y207H mutant was made as a YT-II-100 resistant control, JTE-607 binds in the same active site of CPSF3 so we chose to use this cell line as a control for JTE-607 specificity as well. R-loop formation was not increased in Y207H cells, consistent with a role for CPSF3 in global R-loop formation.

### CPSF3 inhibition has complex impacts on specific R-loop forming loci

Taken together, our data from the use of JTE-607 and YT-II-100 suggest that inhibition of CPSF3 leads to increased R-loop formation throughout the cell. We then tested whether these compounds affect R-loop formation at specific genes that are known to form R-loops and are often used as hallmarks for studies of R-loop regulation and biology. To accomplish this, we evaluated R-loop abundance at specific genes previously reported to be involved in R-loop formation: *RPL13A, TRIM33, CALM3, TFPT, FBXL17*, and *MEPCE*^32,33^. We also evaluated genes used as negative controls for R-loop formation: *EGR1*, and *SNRPN*^32,33^.

We used DNA/RNA immunoprecipitation (DRIP)^32,33^ to isolate R-loops from cell lysate using the S9.6 antibody followed by qPCR to measure abundance of the isolated R-loops at specific genomic loci. As with the dot blots described above, we used the compound resistant mutant HCT116 cell line (Y207H) and pre-treatment with RNase H to gain insights into whether any observed changes in R-loop formation were likely to be on-target effects due to the control of CPSF3 activity by YT-II-100 or JTE-607.

After treating wild-type HCT116 cells with YT-II-100, we observed no increase or non-significant increases in R-loops at the loci known to form R-loops (**Fig. 3A**). Treatment of Y207H mutant HCT116 cells with YT-II-100 also led either to no increase or a non-significant (p=0.1) increases in R-loops at these positive loci (**Fig. 3B**). The failure to observe significant R-loop increases, in combination with the failure to observe a difference between wild-type HCT116 and Y207H HCT116 cells is consistent with the conclusion that YT-II-100 does not affect R-loops at these loci through inhibition of CPSF3. The non-significant increases in R-loop formation at the *RPL13A, TRIM33, CALM3*, and *MEPCE* loci in both HCT116 and Y207H cells hint at YT-II-100 causing increases in R-loop formation that are independent of binding to CPSF3.

**Figure 3.**
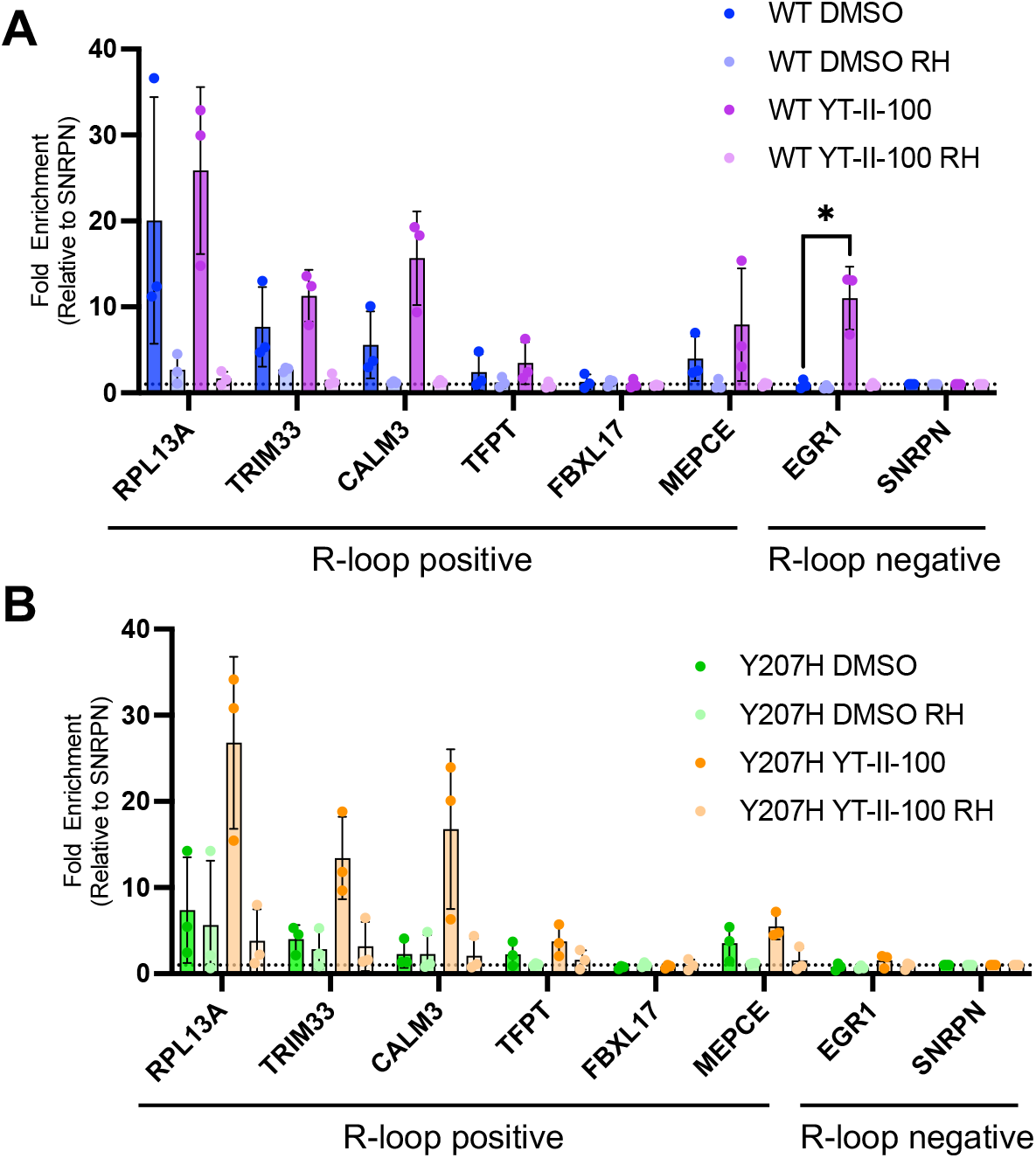
Inhibition of CPSF3 with YT-II-100 has mixed influence on specific R-loop loci. DRIP-qPCR data plotted as fold enrichment relative to negative control gene (SNRPN) of (A) WT and (B) Y207H cells treated with DMSO or YT-II-100 for 6 hours. Samples that were pre-treated with RNase H are denoted as RH. Points are plotted as the average of biological replicates +/-SD. Significance was analyzed by Mann-Whitney test and denoted with *: EGR1 P=0.0286

In the wild-type cells, we observed a significant increase in R-loop detection at one of the negative control loci, the intergenic region downstream of *EGR1* (Fig. 3A). This increase in R-loop abundance was detected in triplicate from the wild-type cells treated with YT-II-100, but not in the DMSO control or in mutant Y207H cells (Fig. 2AB). Because this increase was not present when RNase H was added or in the mutant Y207H cells, it is likely that inhibition of CPSF3 by YT-II-100 plays a role. The second gene that was not known to form R-loops, SNRPN, did not show an increase in R-loop formation.

We also evaluated compound JTE-607 for its ability to control R-loop formation in wild type HCT116 cells (**Fig. 4A**) and mutant Y207H HCT116 (**Fig. 4B**) cells. As we had observed with compound YT-II-100, we observed no changes in R-loop formation within wild-type HCT116 cells or Y207H at the *RPL13A, TRIM33, CAL3, TFPT, FBXL17*, and *MEPCE* genes (Fig. 4AB). Unlike in the YT-II-100 treated samples however, we did not observe an increase at the EGR1 locus in the JTE-607 treated samples (Fig. 4A) or SNRPN, the other negative control gene.

**Figure 4.**
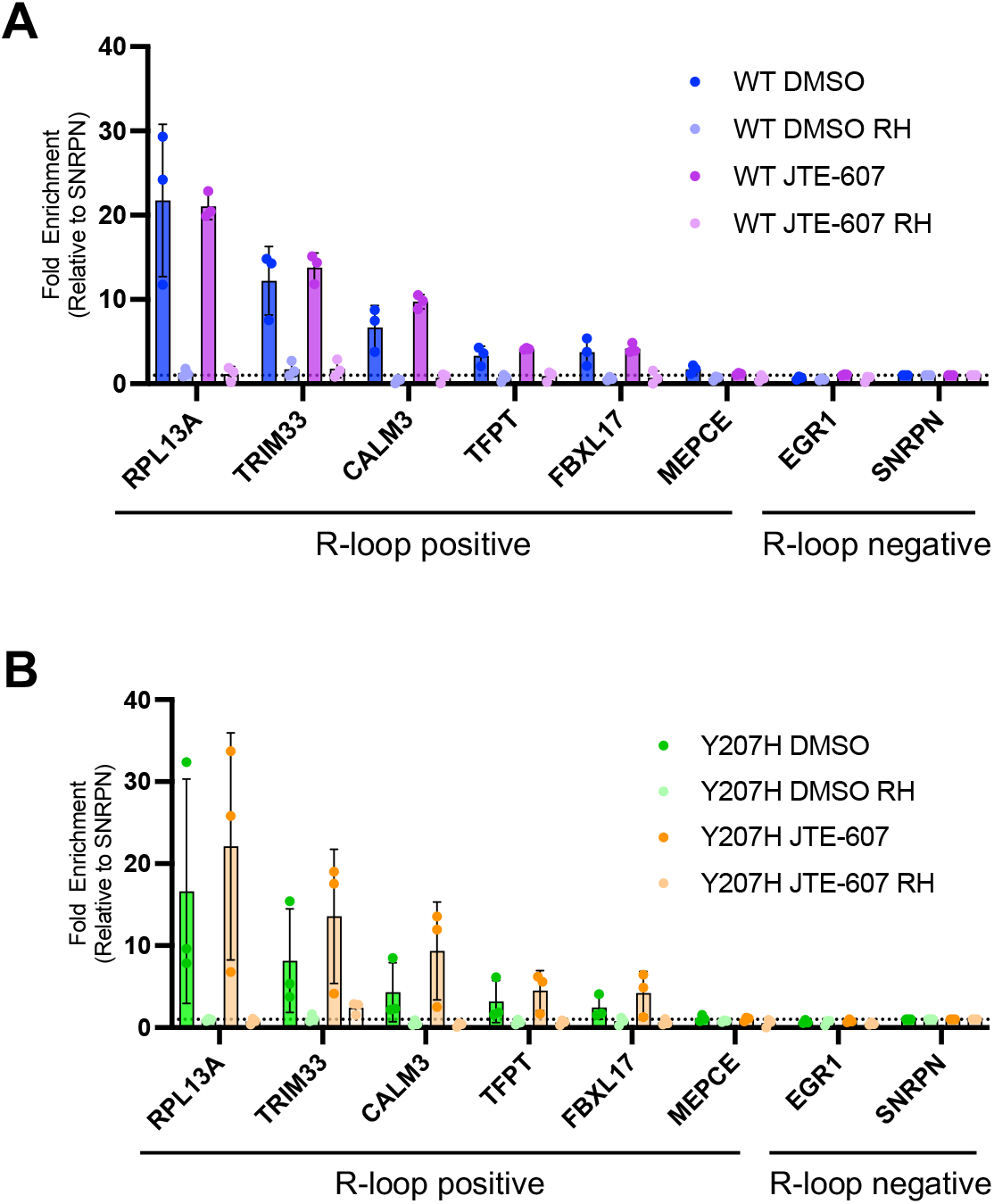
Inhibition of CPSF3 with JTE-607 has mixed influence on specific R-loop loci. DRIP-qPCR data plotted as fold enrichment relative to negative control gene (SNRPN) of (A) WT and (B) Y207H cells treated with DMSO or JTE-607 for 6 hours. Samples that were pre-treated with RNase H are denoted as RH. Points are plotted as the average of biological replicates +/-SD. Significance was analyzed by Mann-Whitney test.

## Discussion

R-loops are biological structures that require tight regulation^9,10,14,32,34^. Their broad roles in transcription, DNA damage repair, and overall genomic stability must be balanced to avoid the negative impacts of untimed or excess formation. Small molecules like YT-II-100 that control R-loop formation in a predictable manner would be important biological probes and possible lead compounds for therapeutic investigation. Previous reports using a different small molecule inhibitor, JTE-607, produced mixed results regarding CPSF3 inhibition and R-loops. In AML and Ewing’s sarcoma cell lines, JTE-607 caused a global increase in R-loops^21^. However, in an ovarian cancer model, JTE-607 was observed to have no significant effect on endogenous levels of R-loops^22^.

Here, we examined the impact of CPSF3 inhibition by YT-II-100 and JTE-607 on R-loop formation and accumulation. We observed global changes in R-loop abundance in a compound-dependent manner when colorectal cancer cells (HCT116) were treated with the novel CPSF3 inhibitor, YT-II-100 (Fig. 1D) or previously described inhibitor JTE-607. To our surprise, however, neither YT-II-100 nor JT-607 produced statistically significant increases R-loop formation at loci that had been previously characterized to form R-loops (Figs. 3 and 4).

Why would the addition of the two different inhibitors of CPSF3 to cells affect global R-loop formation but not the formation of R-loops at hallmark genes that are known to form R-loops and are often used as standards for studies of R-loop formation? A full answer will require subsequent investigation. However, we hypothesize that since these hallmark genes were already forming R-loops, the addition of YT-II-100 or JT-607 will have less opportunity to exert an effect. In these cells, under the growth conditions used in our studies, R-loop formation at the hallmark genes may already be maximal.

Enhanced R-loop formation was observed at one of our negative control genes, the intergenic region downstream of *EGR1*. Our experiments with RNase H and HCT116 Y207H cells are consistent with this being an “on-target” effect due to inhibition of CPSF3 by YT-II-100. Treatment with YT-II-100 blocks proper transcription termination and polyadenylation, leading to transcription readthrough, including at the *EGR1* gene^23^. The formation of an R-loop at *EGR1* could, therefore, be a result of this readthrough producing a transcript that is not present under normal conditions. Alternatively, it is possible that an R-loop already forms at very low levels at this locus, but RNAPII may slow down enough to increase the formation of stable R-loops that can be detected by the S9.6 antibody. Finally, we did not observe increased R-loop formation at EGR1 after using JTE-607, suggesting that the mechanism may be more complex or possibly an off-target effect. Alternatively, JTE-607 is a pro-drug that must be activated by the enzyme CES1^21,35^. It is possible that we are not observing the full activity of this compound in our studies if the expression level or activity level is suboptimal relative to the concentration of compound used.

Our mixed findings are consistent with R-loop formation being a complex phenomenon^34^. Additional work is necessary to provide the mechanistic details necessary to understand why some loci respond to YT-II-100 by increasing R-loop formation while others do not. DNA/RNA immunoprecipitation sequencing (DRIP-seq) uses the S9.6 antibody to globally survey sites of R-loop formation^32,33,36^. DRIP-seq would produce genome-wide, site-specific information about where R-loops form in HCT116 and HCT116 Y207H cells and how R-loop formation changes at these sites when YT-II-100 is added. A catalog of the sites that are sensitive to YT-II-100 would help predict the therapeutic potential of this small molecule.

## Conclusions

Our data support the conclusion that the inhibition of CPSF3 by YT-II-100 and JTE-607 increase the global abundance of R-loops. This increase is not uniform. It does not occur at several loci known to form R-loops but does occur at a locus where R-loops are not normally observed. Genome-wide analysis of R-loops following CPSF3 inhibition, experimental validation, and mechanistic investigation will be necessary to fully understand how genes and R-loop formation are affected by CPSF3 and its inhibition by small molecules. Our work suggests that R-loop formation and the interplay between CPSF3 and its small molecule inhibitors are a complex phenomenon that depends on the context of each gene. Future development of synthetic R-loop regulators will benefit from recognizing this complexity.

## Author contributions

This manuscript was prepared by C.H. and D.R.C. Data was collected and analyzed by C.H., V.T., and J.H. with supervision from D.R.C.

## Acknowledgements

The authors thank Deepak Nijhawan, Ye Tao, and Jef DeBrabander for providing the cell lines and YT-II-100 compound used in this study. DRC was supported by the National Institutes of Health (NIH) (GM118103) and the Robert Welch Foundation (I-2184). DRC holds the Rusty Kelley Professorship in Medical Science.

## Supplementary information

## Materials and Methods

### Reagents

YT-II-100 was a gift from the Nijhawan lab. Synthesis was as previously described^1^. JTE-607 was purchased from Millipore Sigma (SML2833).

### Cell culture

HCT116 (ATCC) and HCT116 Y170H cells were a gift from the Nijhawan lab and used as previously described^1^. Cells were grown in Dulbecco’s Modified Eagle Medium High Glucose (D5796, Sigma) and supplemented with 10% FBS and 10 mM L-glutamine. All cells were cultured at 37^°^C in 5% CO_2_.

### Cell treatment and genomic DNA isolation

HCT116 and HCT116 Y207H were seeded at 5×10^6^ cells per well in 6-well plates. When cells reached ∼80% confluence, cells were treated with either DMSO or 5µM YT-II-100 for 6 hours. Cells were collected with trypsin. Cells were washed with PBS and nuclei were isolated using hypotonic lysis buffer (10 mM Tris HCl pH 7.4, 10 mM NaCl, 3 mM MgCl_2_, 2.5% NP-40, 0.5 mM DTT). 10 million cells were resuspended in 700 µL of TE buffer pH 8 and treated with proteinase K and 1% SDS for 3 hours at 42^°^C. Samples were sonicated, followed by phenol-chloroform extraction and ethanol precipitation.

### Dot Blot

Dot blot was performed as previously described^2^. Briefly, genomic DNA from treated cells (Cell treatment and genomic DNA isolation) was diluted and dotted on nitrocellulose membrane (10600002, Cytvia Amersham) and allowed to dry. Samples were UV crosslinked and membranes were blocked with 5% milk in PBST for 1 hour. Membranes were incubated with S9.6 antibody (MABE1095, Sigma) overnight. Membranes were washed 3X with PBST prior to addition of HRP conjugated secondary antibody (anti-mouse, Bethyl Laboratories). Membranes were washed again 3X with PBST. Chemiluminescent substrate (34577, Thermo Scientific) was added to samples and exposed onto film.

### DNA/RNA immunoprecipitation (DRIP)

DRIP was performed as previously described with modifications^3^. Briefly, cells were treated, and genomic DNA was isolated as in cell treatment and genomic DNA isolation. Half of each sample was pre-treated with RNase H for 6 hours at 37^°^C. Samples with and without RNase H treatment were precleared for 2 hours before addition of S9.6 (MABE1095, Sigma) for overnight binding. RNA/DNA hybrids were recovered with 80µL of protein G Dynabeads, washed twice and eluted. Eluted samples were treated with proteinase K, phenol-chloroform extracted, and ethanol precipitated. Validation of DRIP was performed by qPCR using primers (Supp. Table 1) as previously described^3,4^. Mann-Whitney tests were used for statistical analysis.

### Immunofluorescence microscopy

HCT116 cells were seeded at 1.5×10^5^ cells per well in cover glass (pre-coated with poly-L-lysine) in 24-well plates. After 24 hours, cells were treated with either DMSO or 5µM YT-II-100 for 6 hours. After YT-II-100 treatment, the cells were extracted with 0.1% Triton X in PBS for 1 minute to remove the cytoplasm. Samples were then fixed with 4% formaldehyde in 1X PBS and permeabilized with 0.2% Triton X in PBS. Cells were blocked with 10% NGS for 1 hour and then incubated with S9.6 antibody (1:1000) at 4°C overnight. On the next day, cells were washed three times with PBS and incubated with anti-mouse Alexa Fluor™ Plus 488 antibody (1:1000) for 1 hour at room temperature. The cells were then stained with mounting media with DAPI (Vector Labs, H-1500).

Cells were imaged at 60× magnification using a Widefield Deltavision microscope. Images were processed by blind deconvolution with AutoQuant X3, and finally analyzed using ImageJ/Fiji. For each experiment, images were taken in parallel, with identical exposure times and settings. When the final images were processed by ImageJ/Fiji, the same brightness/contrast parameter was applied for comparison.

---

**Supplementary table 1.**
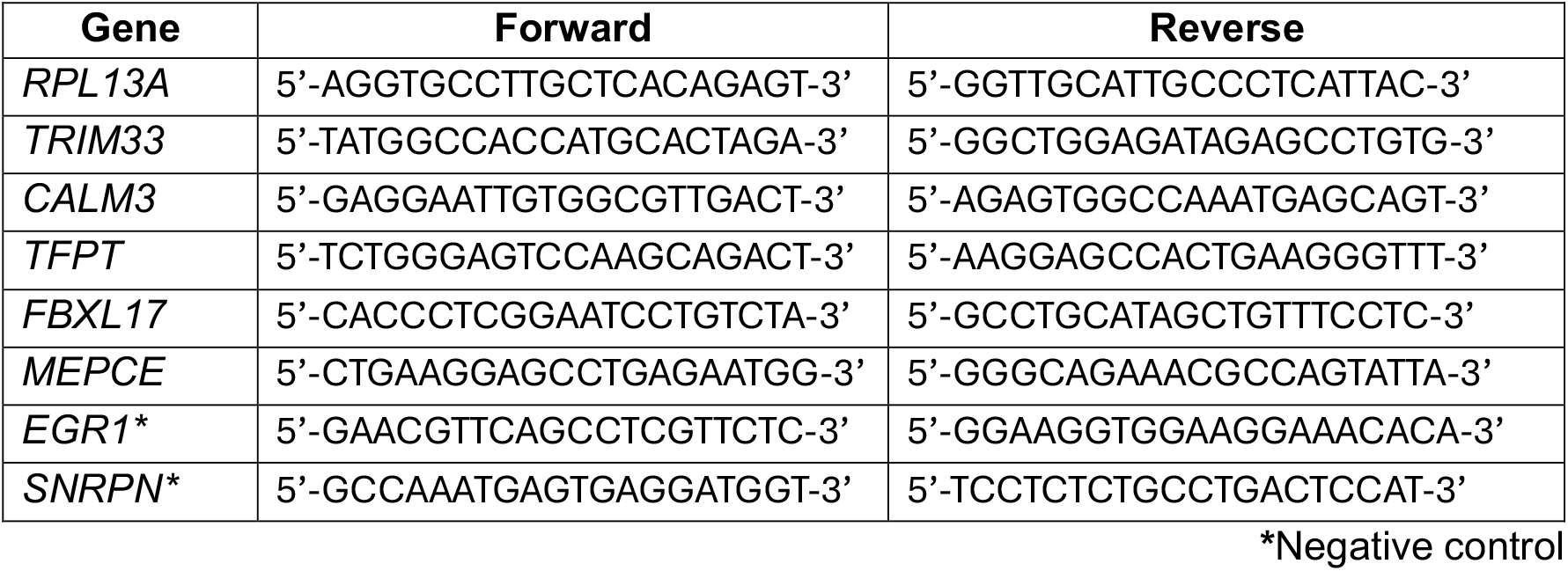
Primer sequences used for DRIP-qPCR.

